# *GmSALT3* expression improves reactive oxygen species detoxification in salt-stressed soybean roots

**DOI:** 10.1101/2021.05.26.445725

**Authors:** Yue Qu, Lili Yu, Rongxia Guan, Oliver Berkowitz, Rakesh David, James Whelan, Melanie Ford, Stefanie Wege, Lijuan Qiu, Matthew Gilliham

## Abstract

Soybean plants are salinity (NaCl) sensitive, with their yield significantly decreased under moderately saline conditions. *GmSALT3* is the dominant gene underlying a major QTL for salt tolerance in soybean. *GmSALT3* encodes a transmembrane protein belonging to the plant cation/proton exchanger (CHX) family. It is currently unknown through which molecular mechanism(s) the ER-localised GmSALT3 contributes to salinity tolerance, as its localisation excludes direct involvement in ion exclusion. In order to gain insights into potential molecular mechanism(s), we used RNA-seq analysis of roots from two soybean NILs (Near Isogenic Lines); NIL-S (salt-sensitive, *Gmsalt3*) and NIL-T (salt-tolerant, *GmSALT3*), grown under control and saline conditions (200 mM NaCl) at three time points (0h, 6h, and 3 days). Gene ontology (GO) analysis showed that NIL-T has greater responses aligned to oxidation reduction. ROS were shown less abundant and scavenging enzyme activity was higher in NIL-T, consistent with the RNA-seq data. Further analysis indicated that genes related to calcium signalling, vesicle trafficking and Casparian strip (CS) development were upregulated in NIL-T following salt treatment. We propose that GmSALT3 improves the ability of NIL-T to cope with saline stress through preventing ROS overaccumulation in roots, and potentially modulating Ca^2+^ signalling, vesicle trafficking and formation of diffusion barriers.

**Highlight:** RNA-seq analysis revealed that GmSALT3, which confers improved salt tolerance on soybean, improves reactive oxygen species detoxification in roots.

## Introduction

Plants use reactive oxygen species (ROS) as signalling molecules at low concentrations to control and regulate various biological processes, such as growth, programmed cell death, hormone signalling and development (Pei *et al*., 2000; Mittler, 2002; Neill *et al*.; 2002, Foreman *et al*., 2003; Overmyer *et al*., 2003). However, ROS are also a toxic by-product of many reactions, which subsequently need to be eliminated by ROS scavenging enzymes. Under non-stressed conditions, the production of ROS and the capacity of the cell to scavenge them are generally at equilibrium (Foyer and Noctor, 2005). However, under biotic and abiotic stresses such as salinity, drought, temperature extremes, flooding, heavy metals, nutrient deprivation, and pathogen attack, intracellular ROS levels can increase dramatically. ROS at high concentrations damage plant cells through lipid peroxidation, DNA damage, protein denaturation, carbohydrate oxidation, and enzymatic activity impairment leading to significant damage to cellular functions and even cell death (Noctor and Foyer, 1998; Mittler *et al*., 2004; Foyer and Noctor, 2005; Gill and Tuteja, 2010). ROS damage due to environmental stress is therefore a leading factor contributing to reduced global crop production. ROS have, for example, detrimental effects on membrane integrity under drought stress and cause early senescence, lowering crop yield (Mittler, 2002; Sharma *et al*., 2017). In plants, ROS are eliminated through enzymatic and non-enzymatic pathways. Enzymatic antioxidant systems include scavenger enzymes such as SOD (superoxide dismutase), APX (ascorbate peroxidase), GPX (Glutathione peroxidases), PrxR (proxiredoxin), GST (glutathione-S-transferase), and CAT (Catalase); while non-enzymatic pathways include low molecular metabolites, such as ASH (ascorbate), GSH (glutathione), proline, α-tocopherol (vitamin E), carotenoids and flavonoids (Mittler *et al*., 2004; Feroza *et al*., 2016).

Among the above-mentioned stresses, salinity is one of the most prominent factors restricting crop production and agricultural economic growth worldwide. More than US$12 billion in revenue is estimated to be lost annually because of saline-affected agricultural land areas (Flowers *et al*., 2010; Bose *et al*., 2014; Gilliham *et al*., 2017). Soybean (*Glycine max* (L.) Merrill) is an important legume crop that contributes 30% of the total vegetable oil consumed and 69% of human food and animal feed protein-rich supplements (Prakash, 2001; Lam *et al*., 2010). The yield of soybean can be significantly reduced by salinity stress, especially during the early vegetative growth stage (Pi *et al*., 2016).

In soybean, *GmSALT3* (also known as *GmCHX1* and *GmNcl*) has been identified as a dominant gene that confers improved salinity tolerance (Guan *et al*., 2014; Qi *et al*., 2014, Do *et al*., 2016). It is mainly expressed in root phloem-and xylem-associated cells and the protein is localized to the ER (endoplasmic reticulum), not the plasma membrane (PM) (Guan *et al*., 2014). NILs (near isogeneic lines), differing only in a transposon insertion in the *GmSALT3* gene, show that *GmSALT3* is involved in shoot exclusion of both, Na^+^ (sodium) and Cl^−^ (chloride). In NIL-Ts with a functional GmSALT3, leaf Cl^−^ exclusion can be observed prior to Na^+^ exclusion (Liu *et al*., 2016), suggesting two distinct mechanisms for the exclusion of the two ions. Grafting of NILs showed that shoot Na^+^ exclusion occurs via a root xylem-based mechanism and shoot Cl^−^ exclusion likely depends upon novel phloem-based Cl^−^ recirculation (Qu *et al*., 2021). Additionally, *GmSALT3* is also related to K^+^ homeostasis (Do *et al*., 2016; Qu *et al*., 2021). Despite the known importance of GmSALT3 for soybean salinity tolerance, the molecular mechanism through which the ER-localised GmSALT3 contributes to improved salinity stress tolerance has not yet been revealed.

There are studies providing a basis for examining the general responses of soybean roots to salt stress, however, these studies compared genetically diverse soybeans that differ in many aspects. High-throughput “-omics” technologies, including transcriptomics, proteomics, and metabolomics have been applied in an attempt to understand soybean root responses to salinity stress (Aghaei *et al*., 2009; Toorchi *et al*., 2009; Ge *et al*., 2010; Lu *et al*., 2013; Qin *et al*., 2013; Pi *et al*., 2016). For instance, proteomic studies utilising the *Glycine max* cultivar Wenfeng07 (relatively salt-tolerant) and the *Glycine soja* (wild soybean) cultivar Union85-140 (relatively salt-sensitive), suggested that Wenfeng07’s tolerance to salinity stress was associated with flavonoid accumulation in the tolerant accession. Flavonoids belong to the metabolites that scavenge ROS, and flavonoid synthesis is dependent on the activity of a few key enzymes, including chalcone synthase (CHS), chalcone isomerase (CHI) and cytochrome P450 monooxygenase (CPM) (Pi *et al*., 2016). Wenfeng07 contains the *GmSALT3* salt tolerant allele and it is absent in Union85-140, but an explicit link between ROS detoxification, GmSALT3 and salt tolerance is not possible in this diverse genetic material.

We took advantage of our previously isolated NILs, and performed RNA-sequencing of NIL-T (salt-tolerant, *GmSALT3*) and NIL-S (salt-sensitive, *Gmsalt3*) roots, to further investigate the mechanism by which GmSALT3 confers salinity tolerance. Genes connected to ROS signalling were significantly differently regulated in NIL-T compared to NIL-S under saline conditions, suggesting a direct connection of ROS production to GmSALT3. Enzymatic assays further revealed that presence of *GmSALT3* is essential for maintaining ROS homeostasis under saline conditions. Differential gene expression under control conditions suggested that NIL-S might have a higher biotic resistance compared to NIL-T.

## Materials and methods

### Plant growth conditions and stress treatments

NIL-T (Salt-tolerant, *GmSALT3*) and NIL-S (Salt-sensitive, *Gmsalt3*) plants were grown in a growth chamber (RXZ-500D; Ningbo Jiangnan Instrument, China), with a day length of 16h (with a light-emitting diode light source at 400 μmol m^−2^ s^−1^) at 28 °C, and 8 h dark at 25 °C, with 60% relative humidity throughout. Soybean seedlings were treated with 200 mM NaCl (salt treatment) or water (control) at 10 days after sowing (DAS). 200 mM NaCl or water were applied again at 12 DAS.

### Total RNA extraction and RNA-seq library construction

Total RNA was extracted from soybean roots using TRIZOL reagent (Ambion, http://www.ambion.com). Root samples were harvested at three time points, 0h, 6h, and 3d of a 200 mM salt-treatment with the corresponding non-treatment controls. To remove the residual DNA, the extracted RNA was treated with RNase-free DNase I (New England Biolabs, https://www.neb.com) for 30 min at 37°C. Thirty RNA libraries were generated for paired-end reads using an Illumina HiSeq 2500 sequencer, consisting of 3 biological samples per time point per genotype.

### RNA-seq data analysis and assembly

Raw reads were generated and mapped to the latest soybean genome sequence Gmax_275 Wm82.a2.v1 (Glyma 2.0) using TopHat2 (Kim *et al*., 2013). Clean mapped reads were obtained by removing low quality (Q<30) sequences, adapter fragments and barcode sequences. A quality control test by fastQC (Andrews, 2010) revealed the quality of RNA-seq libraries construction and sequence alignment were sufficient for further analysis.

### Gene expression level and DEG analysis

Gene expression was compared in FPKMs (Fragments Per Kilobase of transcript per Million) calculated using the Cufflinks functions cuffquant and cuffnorm (Trapnell *et al*., 2012). Differential expression analysis of two conditions/groups was performed using the DESeq R package (1.10.1). The resulting P values were adjusted using the Benjamini and Hochberg’s approach for controlling the false discovery rate. The cut-offs for differentially expressed genes (DEGs) were Log_2_FC (fold change) ≥ 1, FDR (False Discover Rate) < 0.01.

### GO enrichment and KEGG pathway analysis

GO enrichment analysis of DEGs was implemented using AgriGO (Du *et al*., 2010). GO terms with corrected *p* values < 0.05 were deemed significantly enriched. KEGG (Kanehisa *et al*., 2007) is a database resource for understanding high-level functions and utilities of the biological system, such as the cell, the organism and the ecosystem, from molecular-level information, especially large-scale molecular datasets generated by genome sequencing and other high-throughput experimental technologies (http://www.genome.jp/kegg/). We used KOBAS (Mao *et al*., 2005) software to test the statistical enrichment of differential expression genes in KEGG pathways.

### Weighted Gene Co-expression Network Analysis (WGCNA)

There were 54,175 genes annotated in all the RNA-seq libraries. After filtering out non-expressed genes, low-expressed genes (FPKM < 2) and genes that possessed large expression variation across replicates, 28,223 genes were selected for WGCNA analysis. An unsigned gene co-expression network was built using the WGCNA R-package (Langfelder and Horvath, 2008). The Topological Overlap Measure (TOM) was calculated using the adjacency matrix between all genes with a power of 18. A gene dendrogram was created by using the dissimilarity TOM. Branches of the dendrogram (Supplementary Fig. 6) indicated modules (clustered highly co-expressed genes) through using the Dynamic Tree Cut algorithm (Langfelder et al., 2008). Each module has an assigned colour. Similar modules were merged by calculating module eigengenes. Using the module eigengenes, the Module – Trait Relationships (MTRs) were plotted by calculating the Pearson’s correlations between the module eigengene and the samples (traits). A correlation of 0.75 with at least one of the selected traits was the criteria used to generate modules. Clustered genes in modules of interest were then used for further analysis. Genes annotation and GO term analysis were performed in Soybase (Grant *et al*., 2010) and AgriGO (Du *et al*., 2010), respectively.

### ROS production and contents measurement

Fourteen-day-old NIL-T and NIL-S soybean seedlings were used to measure the generation of ROS in roots. Roots were labelled with 30 μM Dichlorofluorescein diacetate (DCFDA; Sigma-Aldrich, USA), by incubating washed soybean roots for 10 min at room temperature in the dark with 30 μM DCFDA. ROS sensitive fluorescent dyes were imaged using a Nikon SMZ25 stereo microscope (Nikon, Japan). The excitation and emission wavelengths were 488 nm and 500/535 nm, respectively.

ROS contents in soybean root samples were measured using Amplex^®^ UltraRed reagent (Invitrogen, USA). Sodium phosphate buffer (0.5 ml 50 mM pH 7.4) was added to 0.1g soybean powder, after mixing and solubilisation, samples were set on ice for 5 min. Then tubes were centrifuged at 12 x 1000 g for 20 min. 500 μl supernatant was transferred into new tubes. An equal volume of 2:1 (v/v) chloroform : methanol was added and centrifuged at 12 x 1000 g for 5 min. 50 μl aqueous phase was took from each sample and add into each well of 96-well microplate. 50 μl working solution (freshly made) was added into each well. Plates were incubated at 25 °C for 30 min (protected from light). Fluorescence was measured at 540/590 nm.

### Scavenging activity of the superoxide anion (O_2_^-^) assay

The scavenging activity assay was adapted from Pi *et al*. (2016) with slight modifications. Less root homogenate (0.1 g) was used for measurement. Antioxidant enzymes were extracted with 2 ml of 0.05 M phosphate buffer (pH5.5) from 0.1 g root homogenate. The extract was then centrifuged at 12,000 x g (4 °C) for 10 min. Supernatant (40 μl) was added into 160 μl reaction buffer, which contains 80 μl phosphate buffer, 40 μl 0.05 M guaiacol (Sigma, USA), 40 μl 2% hydrogen peroxide (H_2_O_2_). The increased absorbance at 470 nm in 96-well plates due to the enzyme-dependent guaiacol oxidation was recorded every 30s until 4 min of reaction.

### Real-time quantitative reverse-transcription polymerase chain reaction (RT-qPCR) validation for RNA-seq results

Total RNA was extracted from soybean root tissues using TRIZOL reagent (Ambion, http://www.ambion.com). To remove the residual DNA, the extracted RNA was treated with RNase-free DNase I (New England Biolabs, https://www.neb.com) for 30 min at 37°C. For gene expression, first-strand cDNA synthesis was done with a PrimeScript RT Reagent Kit (TaKaRa, Japan, http://www.takara.co.jp/english). Real-time PCR was performed using SYBR Premix Ex Taq II (TliRNaseH Plus) (TaKaRa). The level of *GmSALT3* transcript was normalised using the control gene *GmUKN1* (Hu *et al*., 2009).

## Results

### RNA-sequencing preparation and profiles

*GmSALT3* has previously been shown to be expressed in roots, and grafting experiments have shown that presence of GmSALT3 expression in roots is sufficient to confer shoot Na^+^ and Cl^−^ exclusion (Guan *et al*., 2014; Qu *et al*., 2021). Total root RNA was extracted from NIL-T and NIL-S, and transcriptomes obtained using RNA-seq in a time course from plants grown under control and saline conditions. To investigate short-and long-term responses, root samples were harvested at three time points, 0h (control), 6h, and 3d with 200 mM salt-treatment and controls (Fig. 1a; 1b) and in biological triplicates. Thirty RNA libraries were generated for paired-end reads using an Illumina HiSeq 1500 sequencer. In total, 1.6 billion paired 100 bp raw reads were generated and mapped to the soybean genome sequence Gmax_275 Wm82.a2.v1 (Glyma 2.0) using TopHat2 (Kim *et al*., 2013). The average mapping percentage was 81.25%. After trimming of low quality (Q<30), adapter fragments and barcode sequences, a total of 804 million clean mapped reads were identified. Combined with a quality control test using fastQC (Andrews, 2010), the quality of RNA-seq libraries construction and sequence alignment was deemed sufficient for further analysis. A summary of mapped reads and quality of sequencing is shown in Table 1. Transcripts corresponding to *GmSALT3/Gmsalt3* in the NIL-T and NIL-S in all samples were confirmed. NIL-T transcriptomes contained reads across the entire full-length transcript for *GmSALT3* were obtained, while in NIL-S reads only mapped to the predicted truncated version *Gmsalt3* (Fig. 1c). A flowchart of data analysis is shown in Fig. 1d.

**Table 1.** Reads mapping and quality of sequencing for 30 *Glycine max* root samples.

| Group | Sample | Total Reads | Mapped Reads | Clean reads | Clean bases | GC Content | %≥Q30 |
| --- | --- | --- | --- | --- | --- | --- | --- |
| g1 | Control 0h T1 | 57,699,406 | 49,137,608 (85.16%) | 28,849,703 | 7,212,425,750 | 50.56% | 92.05% |
|  | Control 0h T2 | 59,769,628 | 51,025,677 (85.37%) | 29,884,814 | 7,471,203,500 | 52.37% | 93.30% |
|  | Control 0h T3 | 62,070,060 | 53,862,681 (86.78%) | 31,035,030 | 7,758,757,500 | 50.01% | 93.35% |
| g2 | Control 0h S1 | 43,642,298 | 32,134,039 (73.63%) | 21,821,149 | 5,455,287,250 | 48.70% | 88.44% |
|  | Control 0h S2 | 46,423,344 | 39,574,288 (85.25%) | 23,211,672 | 5,802,918,000 | 49.23% | 94.08% |
|  | Control 0h S3 | 56,017,714 | 43,638,938 (77.90%) | 28,008,857 | 7,002,214,250 | 50.34% | 93.88% |
| g3 | Control 3d T1 | 48,501,908 | 42,642,939 (87.92%) | 24,250,954 | 6,062,738,500 | 48.90% | 93.69% |
|  | Control 3d T2 | 60,432,732 | 53,220,906 (88.07%) | 30,216,366 | 7,554,091,500 | 50.86% | 94.28% |
|  | Control 3d T3 | 48,809,732 | 41,680,164 (85.39%) | 24,404,866 | 6,101,216,500 | 48.29% | 92.51% |
| g4 | Control 3d S1 | 47,058,528 | 39,937,443 (84.87%) | 23,529,264 | 5,882,316,000 | 48.39% | 93.58% |
|  | Control 3d S2 | 48,461,144 | 40,928,669 (84.46%) | 24,230,572 | 6,057,643,000 | 47.83% | 94.18% |
|  | Control 3d S3 | 49,796,814 | 40,986,303 (82.31%) | 24,898,407 | 6,224,601,750 | 50.35% | 94.30% |
| g5 | Control 6h T1 | 51,648,018 | 30,700,905 (59.44%) | 25,824,009 | 6,456,002,250 | 48.37% | 84.72% |
|  | Control 6h T2 | 34,427,522 | 21,774,315 (63.25%) | 17,213,761 | 4,303,440,250 | 48.21% | 87.04% |
|  | Control 6h T3 | 84,627,144 | 55,643,274 (65.75%) | 42,313,572 | 10,578,393,000 | 48.00% | 86.74% |
| g6 | Control 6h S1 | 62,158,510 | 50,330,267 (80.97%) | 31,079,255 | 7,769,813,750 | 48.61% | 92.06% |
|  | Control 6h S2 | 59,768,732 | 44,180,997 (73.92%) | 29,884,366 | 7,471,091,500 | 48.12% | 89.44% |
|  | Control 6h S3 | 48,852,622 | 41,445,404 (84.84%) | 24,426,311 | 6,106,577,750 | 49.05% | 93.76% |
| g7 | NaCl 3d T1 | 47,478,210 | 39,318,129 (82.81%) | 23,739,105 | 5,934,776,250 | 49.12% | 93.71% |
|  | NaCl 3d T2 | 49,524,852 | 42,007,601 (84.82%) | 24,762,426 | 6,190,606,500 | 50.44% | 94.94% |
|  | NaCl 3d T3 | 53,918,342 | 45,880,121 (85.09%) | 26,959,171 | 6,739,792,750 | 52.93% | 93.81% |
| g8 | NaCl 3d S1 | 55,228,034 | 45,771,588 (82.88%) | 27,614,017 | 6,903,504,250 | 47.24% | 92.28% |
|  | NaCl 3d S2 | 57,398,436 | 46,291,922 (80.65%) | 28,699,218 | 7,174,804,500 | 48.58% | 92.30% |
|  | NaCl 3d S3 | 58,773,304 | 47,497,496 (80.81%) | 29,386,652 | 7,346,663,000 | 48.48% | 91.31% |
| g9 | NaCl 6h T1 | 53,174,654 | 45,735,099 (86.01%) | 26,587,327 | 6,646,831,750 | 51.47% | 93.56% |
|  | NaCl 6h T2 | 52,045,484 | 43,779,481 (84.12%) | 26,022,742 | 6,505,685,500 | 50.75% | 93.30% |
|  | NaCl 6h T3 | 55,579,826 | 45,167,976 (81.27%) | 27,789,913 | 6,947,478,250 | 51.54% | 92.62% |
| g10 | NaCl 6h S1 | 54,197,662 | 45,095,587 (83.21%) | 27,098,831 | 6,774,707,750 | 48.34% | 92.64% |
|  | NaCl 6h S2 | 53,274,670 | 45,180,893 (84.81%) | 26,637,335 | 6,659,333,750 | 50.44% | 92.49% |
|  | NaCl 6h S3 | 47,601,546 | 40,826,998 (85.77%) | 23,800,773 | 5,950,193,250 | 48.16% | 94.75% |

**Fig. 1.**
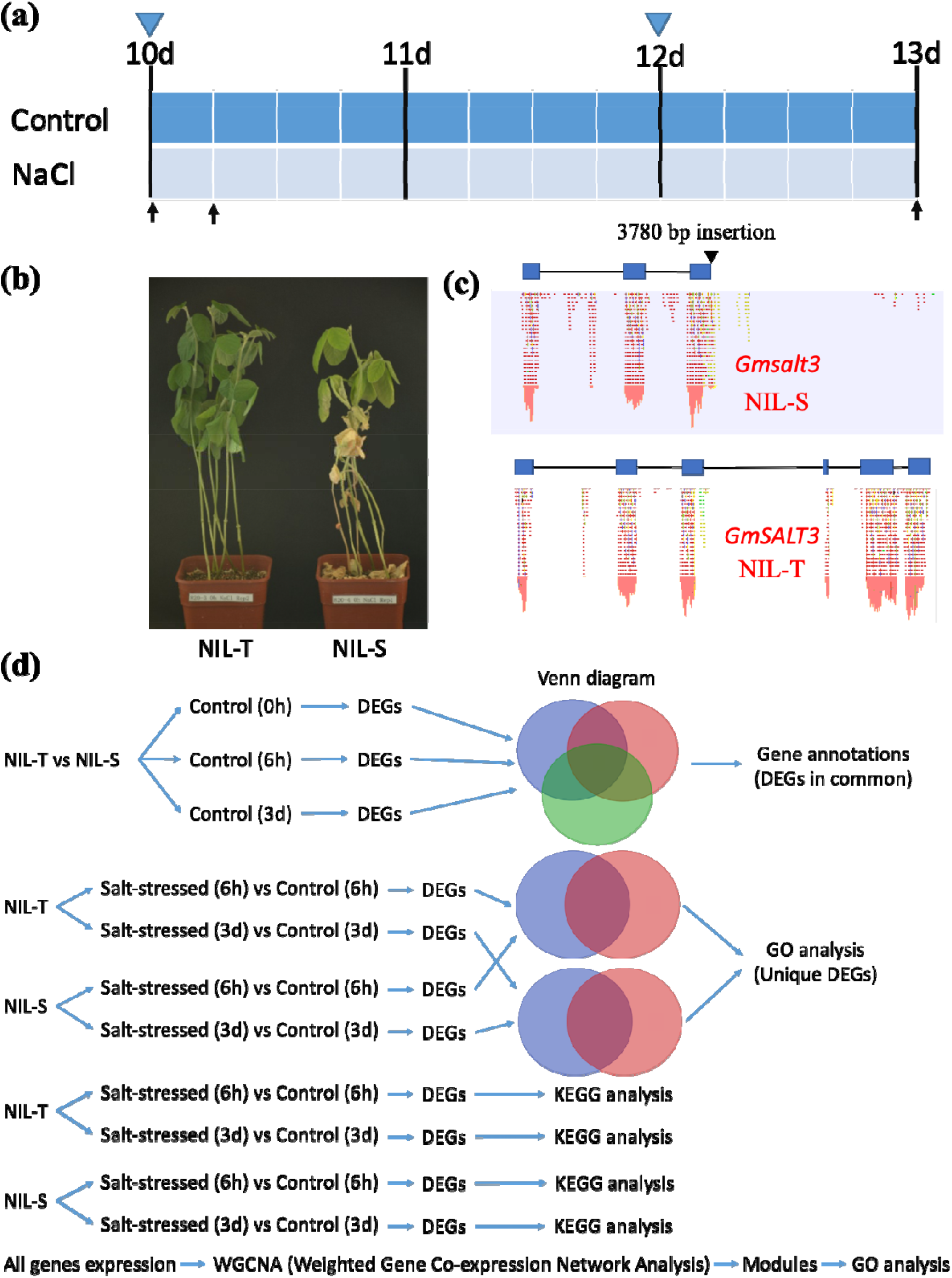
Treatment, sampling strategy and data analysis. (a) Soybean seedlings were treated with 200 mM NaCl (salt treatment) or water (control) at 10 days after sowing (DAS). Arrows mean the sampling time points (0h, 6h, 3d), triangle indicated the time point when water or NaCl solution was applied. (b) Pictures of soybean that had been treated with NaCl solution for 11 days. (c) Mapped-reads to *GmSALT3* (bottom) and *Gmsalt3* (above). (d) Flowchart of data analysis. NIL-T, NIL with *GmSALT3* allele; NIL-S, NIL with *Gmsalt3* allele; DEGs, differentially expressed genes; GO, gene ontology.

### Overview of DEGs between NIL-T and NIL-S under control conditions

Previous work had shown that *GmSALT3* is expressed under both control and saline conditions, suggesting that there might already be a difference in other gene expression between NIL-T and NIL-S under control conditions. We therefore first compared the transcriptomes of NIL-T and NIL-S (indicated as “T” and “S”, respectively) under control conditions, before analysing changes due to salinity treatment. A PCA plot (for the first two principal components) of ten grouped samples shows a good separation between all comparisons, confirming replicates clustered together consistently. Those comparisons are: Control 0h T vs Control 0h S (grey), Control 3d T vs Control 3d S (purple), and Control 6h T vs Control 6h S (brown) (Fig. 2a). There are 5 up-regulated Differentially Expressed Genes (DEGs) at 0h, 9 up-regulated DEGs at 6h, and 6 up-regulated DEGs at 3d in NIL-S (compared to NIL-T), and a Venn diagram demonstrates 3 of these genes are common across three time points. There were 1, 5, and 5 down-regulated DEGs at 0h, 6h, and 3d, respectively, but the identity of the down-regulated DEGs were different between each of the time point (Fig. 2b). The three consistently up-regulated DEGs in NIL-S at all time points under control conditions were *Glyma*.*07G196800* (Linoleate 13S-lipoxygenase 3-1), *Glyma*.*10G143600* (uncharacterized protein), and *Glyma*.*20G105500* (3-hydroxybenzoate 6-hydroxylase 1-like) (Fig. 2c). Fig. 2d shows the expression profile (in FPKM, Fragments Per Kilobase of transcript per Million) of these three genes at all time points of NIL-T and NIL-S under control conditions.

**Fig. 2.**
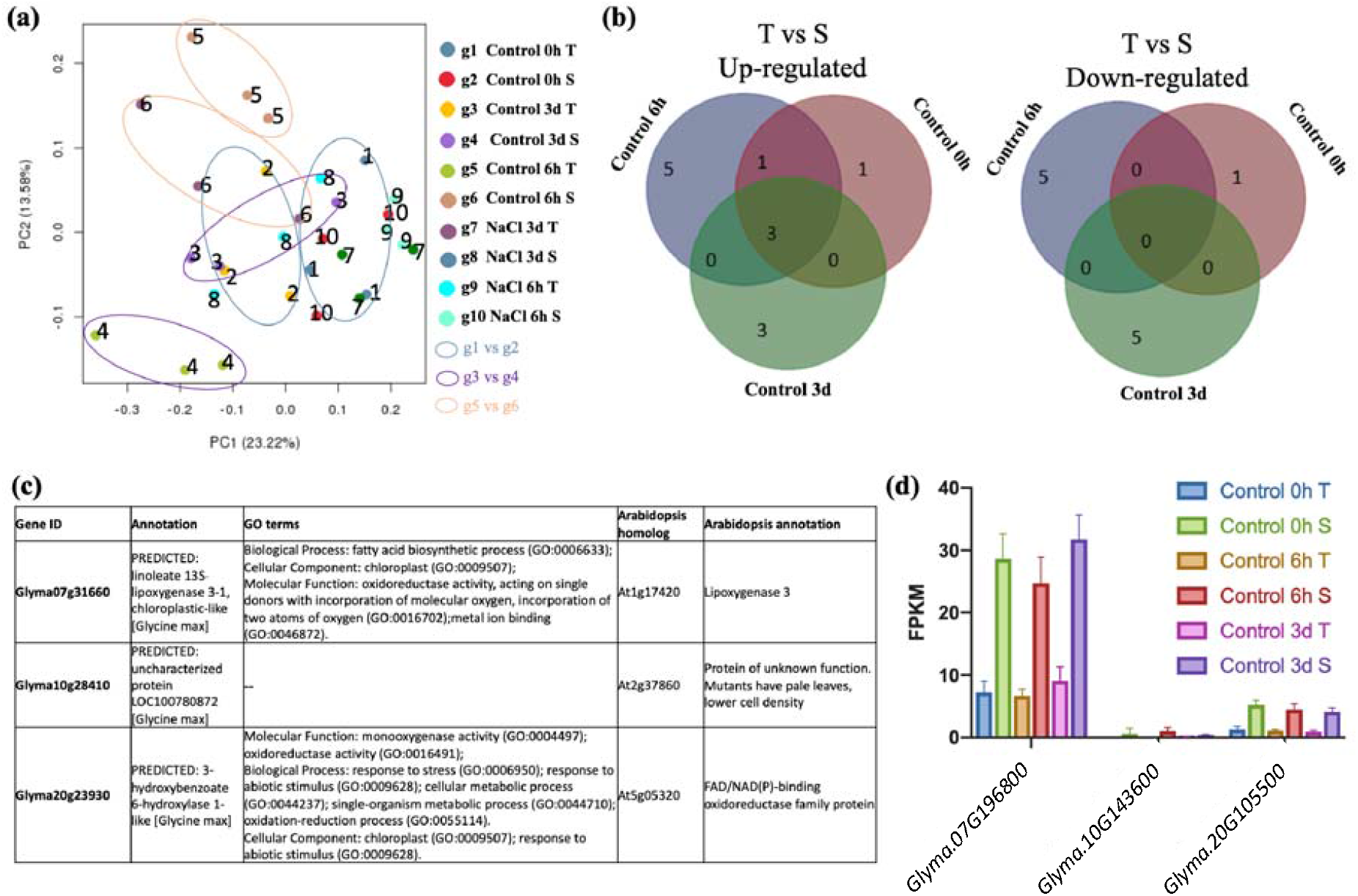
Overview of DEGs between NIL-T salt-tolerant and NIL-S salt-sensitive soybean samples at the 0h, 6h, and 3d timepoints under non-saline conditions. (a) PCA plot of ten groups (g1 – g10). Different coloured dots represent replicates in each group, different coloured circles represent comparisons between groups (g1 vs g2, g3 vs g4, g5 vs g6). (b) Venn diagram of up-and down-regulated DEGs in NIL-S soybeans at 0h, 6h, and 3d under control conditions. (c) Three up-regulated DEGs in NIL-S soybeans at all 0h, 6h, and 3d under control conditions, and their expression profile is shown in (d). The cutoffs for DEGs are Log_2_FC (fold change) ≥ 1, FDR (False Discover Rate) < 0.01.

### Overview of DEGs in salt treated NIL-T and NIL-S when compared to their respective controls

Transcript abundance was compared in FPKMs calculated using the Cufflinks functions cuffquant and cuffnorm (Trapnell *et al*., 2012). DEGs were classified as genes with a Log_2_FC (fold change) ≥ 1 and a FDR (False Discover Rate) < 0.01 when comparing between two conditions. Gene expression differences in response to salt of each genotype was examined separately; to identify the DEGs in NIL-T and in NIL-S in response to salt. Using these parameters and comparing control to salt treated NIL-T, we identified 1816 DEGs that were differently expressed in NIL-T roots in response to salt (1263 up-regulated and 553 down-regulated, 6h T).

We then did the same analysis for NIL-S roots, and compared expression of control and salt treated NIL-S. 6h salt treated NIL-S roots (when compared to control NIL-S) showed a greater initial response in term of DEGs than we found in NIL-T, with a total of 3054 DEGs (1911 up-regulated and 1143 down-regulated, 6h S) (Fig. 3b). After three days of salt treatment, the increased number of DEGs in NIL-S was not observed anymore. On the contrary, the NIL-T now showed a slightly higher number of DEGs compared to control conditions. Now, the NIL-T had 2844 DEGs (1333 up-regulated and 1511 down-regulated, 3d T) while there were 2573 DEGs (1318 up-regulated and 1255 down-regulated, 3d S) in NIL-S roots (Fig. 3b).

**Fig. 3.**
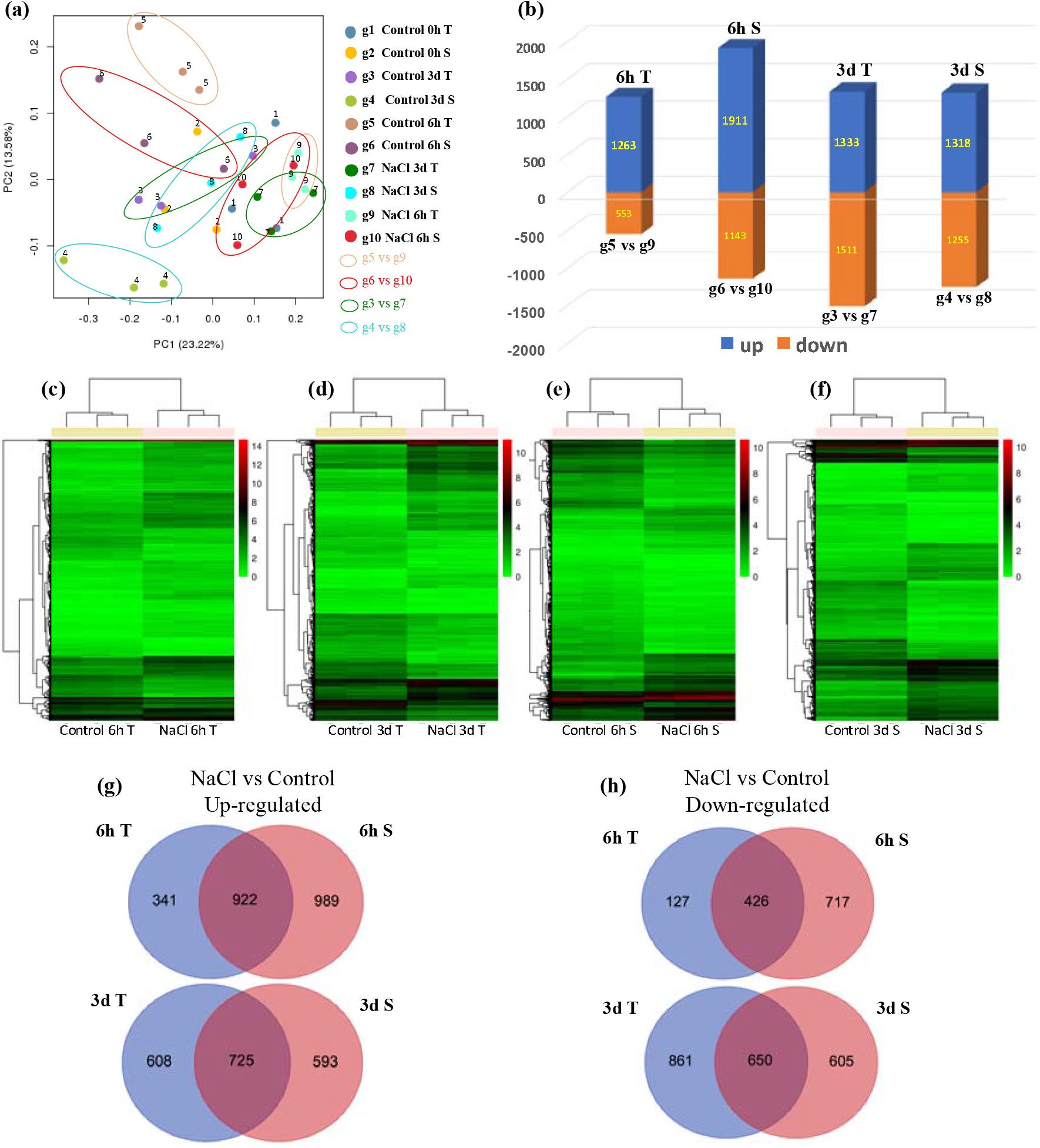
Overview of DEGs (differentially expressed genes) between salt-stressed and control samples in NIL-T and NIL-S soybeans at 6h and 3d. (a) PCA plot of ten groups (g1 – g10). Different coloured dots represent replicates in each group, different coloured circles represent comparisons between groups (g5 vs g9, g6 vs g10, g3 vs g7, g4 vs g8). (b) Numbers of DEGs (up-and down-regulated) between groups. Clustering of DEGs in log_2_FPKMs (Fragments Per Kilobase of transcript per Million) of group 3 to 10 (c-f). The false colour scale from green through to red indicates increasing log_2_FPKM. (g) Venn diagram of up-regulated DEGs in NIL-T and NIL-S soybeans at 6h and 3d under NaCl treatment. (h) Venn diagram of down-regulated DEGs in NIL-T and NIL-S soybeans at 6h and 3d under NaCl treatment. The cut-offs for DEGs are Log_2_FC (fold change) ≥ 1, FDR (False Discover Rate) < 0.01.

Hierarchical clustering of the DEGs of the three replicates in different grouped samples is shown in Fig. 3c-f. These DEGs represent the genes that each soybean line up-or downregulates in response to salt. We then used the hierarchical clustering to identify DEGs, which either similarly change in expression in the two genotypes in response to salt, and which genes show a different change in expression comparing the two genotypes. Analyses of the similarities and differences between the tolerant and sensitive lines indicated that 341 and 989 DEGs are uniquely up-regulated after 6h salt treatment in NIL-T or NIL-S, respectively; and 608 and 593 in the 3d salt treatment, respectively (Fig. 3g). We found similar values for uniquely down-regulated DEGs (Fig. 3h).

### GO analysis of DEGs between control and salt-treated samples of NIL-T and NIL-S

To gain insight into the putative functions of the uniquely up-and down-regulated DEGs at 6h salt treatment, the identified genes were subjected to GO (Gene Ontology) enrichment analysis, the representative GO terms are shown in Table 2, and detailed GO terms are shown in Supplementary Fig. 1 -4.

**Table 2.** GO term analysis of uniquely up-and down-regulated genes under salt treatment in soybean roots at 6h. (a) Up-regulated genes in NIL-T soybean roots. (b) Up-regulated genes in NIL-S soybean roots. (c) Down-regulated genes in NIL-T soybean roots. (d) Down-regulated genes in NIL-S soybean roots. CC, Cellular Component; BP, Biological Process; MF, Molecular Function.

| GO term | Ontology | Description | Number in input list | Number in BG/Ref | p-value | FDR |
| --- | --- | --- | --- | --- | --- | --- |
| GO:0009607 | BP | response to biotic stimulus | 7 | 64 | 9.3e-08 | 2.4e-05 |
| GO:0006952 | BP | defense response | 7 | 118 | 4.4e-06 | 0.00056 |
| GO:0055114 | BP | oxidation reduction | 28 | 2408 | 6.1e-05 | 0.0051 |
| GO:0005507 | MF | copper ion binding | 9 | 219 | 3.4e-06 | 0.00071 |
| GO:0016491 | MF | oxidoreductase activity | 32 | 2744 | 1.5e-05 | 0.0016 |
| GO:0004866 | MF | endopeptidase inhibitor activity | 5 | 92 | 0.00016 | 0.0083 |

(b)
| GO term | Ontology | Description | Number in input list | Number in BG/Ref | p-value | FDR |
| --- | --- | --- | --- | --- | --- | --- |
| GO:0006355 | BP | regulation of transcription, DNA-dependent | 69 | 1874 | 2e-05 | 0.0016 |

(c)
| GO term | Ontology | Description | Number in input list | Number in BG/Ref | p-value | FDR |
| --- | --- | --- | --- | --- | --- | --- |
| GO:0016021 | CC | integral to membrane | 11 | 1419 | 0.00065 | 0.0058 |

(d)
| GO term | Ontology | Description | Number in input list | Number in BG/Ref | p-value | FDR |
| --- | --- | --- | --- | --- | --- | --- |
| GO:0020037 | MF | heme binding | 23 | 728 | 8e-05 | 0.012 |
| GO:0009055 | MF | electron carrier activity | 21 | 702 | 0.00033 | 0.018 |
| GO:0004190 | MF | aspartic-type endopeptidase activity | 9 | 169 | 0.00042 | 0.018 |
| GO:0015035 | MF | protein disulfide oxidoreductase activity | 7 | 96 | 0.00031 | 0.018 |

The enriched GO terms for the 6h T (NIL-T) up-regulated DEGs were all related to stress response such as “oxidation reduction”, “defence response” and “response to biotic stimulus” (Table 2a). Down-regulated GO terms in NIL-T were less specific and mainly associated with “integral to membrane” (Table 2c). For 6h S (NIL-S), a large number of up-regulated genes were associated with the very broad GO term “regulation of transcription, DNA-dependent (GO:0006355)”, and no further downstream classification was considerably enriched (Table 2b). Down-regulated GO terms in NIL-S included “heme binding”, “electron carrier activity”, “aspartic-type endopeptidase activity”, and “protein disulphide oxidoreductase activity” (Table 2d), relating to protein types instead of specific biological process GO terms.

After 3 days of salt treatment, GO analysis of uniquely up-and down-regulated DEGs showed that only “oxidation reduction” and “oxidoreductase activity” were up-regulated in NIL-T (Table 3a). “Oxidation reduction” and “oxidoreductase activity” were consistently up-regulated in NIL-T at both time points. Table 4 shows all the 53 up-regulated genes in “oxidation reduction” and “oxidoreductase activity” GO terms after 3d treatment in NIL-T roots. A group of Cytochrome P450 enzymes-encoding genes were significantly more highly expressed in NIL-T, especially *Glyma*.*13G173500*, which had a FPKM of 279 compared to 71 under control conditions (Table 4). Other genes encoding oxidoreductase enzymes were also included such as peroxidases and dehydrogenases (Table 4). All the gene annotations in each GO are shown in Supplementary Table 1.

**Table 3.** GO term analysis of uniquely up-and down-regulated genes under salt treatment in soybean roots at 3d. (a) Up-regulated genes in NIL-T soybean roots. (b) Up-regulated genes in NIL-S soybean roots. (c) Down-regulated genes in NIL-T soybean roots. (d) Down-regulated genes in NIL-S soybean roots. CC, Cellular Component; BP, Biological Process; MF, Molecular Function.

| GO term | Ontology | Description | Number in input list | Number in BG/Ref | p-value | FDR |
| --- | --- | --- | --- | --- | --- | --- |
| GO:0055114 | BP | oxidation reduction | 53 | 2408 | 8.9e-06 | 0.0037 |
| GO:0016491 | MF | oxidoreductase activity | 57 | 2744 | 2e-05 | 0.0073 |

(b)
| GO term | Ontology | Description | Number in input list | Number in BG/Ref | p-value | FDR |
| --- | --- | --- | --- | --- | --- | --- |
| GO:0006355 | BP | regulation of transcription, DNA-dependent | 51 | 1874 | 9.8e-08 | 3.6e-06 |
| GO:0006468 | BP | protein amino acid phosphorylation | 45 | 2356 | 0.002 | 0.023 |
| GO:0048544 | BP | recognition of pollen | 14 | 135 | 3.8e-09 | 3e-07 |
| GO:0015743 | BP | malate transport | 5 | 34 | 0.0001 | 0.0013 |

(c)
| GO term | Ontology | Description | Number in input list | Number in BG/Ref | p-value | FDR |
| --- | --- | --- | --- | --- | --- | --- |
| GO:0007018 | BP | microtubule-based movement | 17 | 155 | 9.8e-09 | 5.2e-06 |
| GO:0003777 | MF | microtubule motor activity | 17 | 155 | 9.8e-09 | 2.4e-06 |
| GO:0043531 | MF | ADP binding | 27 | 467 | 2.1e-07 | 2e-05 |
| GO:0004565 | MF | beta-galactosidase activity | 6 | 37 | 9.5e-05 | 0.0031 |
| GO:0017171 | MF | serine hydrolase activity | 17 | 330 | 0.00014 | 0.0041 |
| GO:0009341 | CC | beta-galactosidase complex | 6 | 37 | 9.5e-05 | 0.009 |

(d)
| GO term | Ontology | Description | Number in input list | Number in BG/Ref | p-value | FDR |
| --- | --- | --- | --- | --- | --- | --- |
| GO:0055085 | BP | transmembrane transport | 29 | 1167 | 1.9e-05 | 0.0075 |
| GO:0022857 | MF | transmembrane transporter activity | 24 | 1020 | 0.00022 | 0.036 |

**Table 4.**
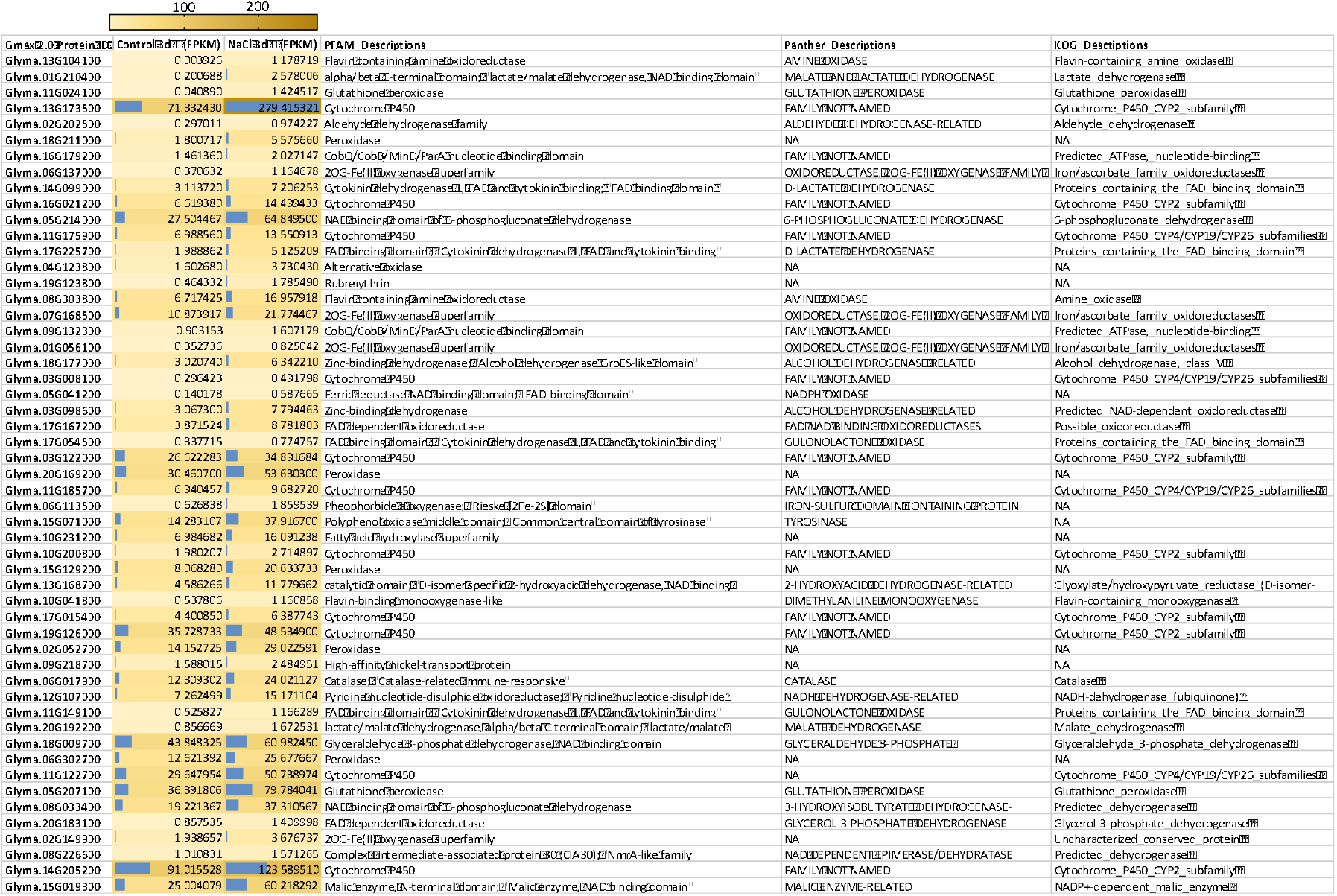
Up-regulated genes in oxidation reduction and oxidoreductase activity GO terms of 3 days salt-treated NIL-T roots. Annotations are achieved from https://www.soybase.org/genomeannotation/.

### KEGG analysis of DEGs between control and salt-treated samples

To understand what pathways were differently altered in NIL-T and NIL-S under salt stress, KEGG (Kyoto Encyclopedia of Genes and Genomes) analysis was performed. This analysis can reveal enrichments of biological pathways, similar to GO term analysis but KEGG takes fold changes of DEGs into account, while the GO term analysis does not.

In our analysis, 1816 and 3054 DEGs in NIL-T and NIL-S, respectively, were enriched in 58 pathways (corrected *p*-Value <0.05) after 6h salt-treatment (Fig. 4a). Most of the 58 pathways in both NIL-T and NIL-S were within the metabolism category with “Phenylpropanoid biosynthesis”, “Phenylalanine metabolism”, and “Starch and sucrose metabolism” as the top three pathways (Fig. 4a). Overall, NIL-S has more DEGs compared to NIL-T after salt-treatment, across all significantly changed pathways.

**Fig. 4.**
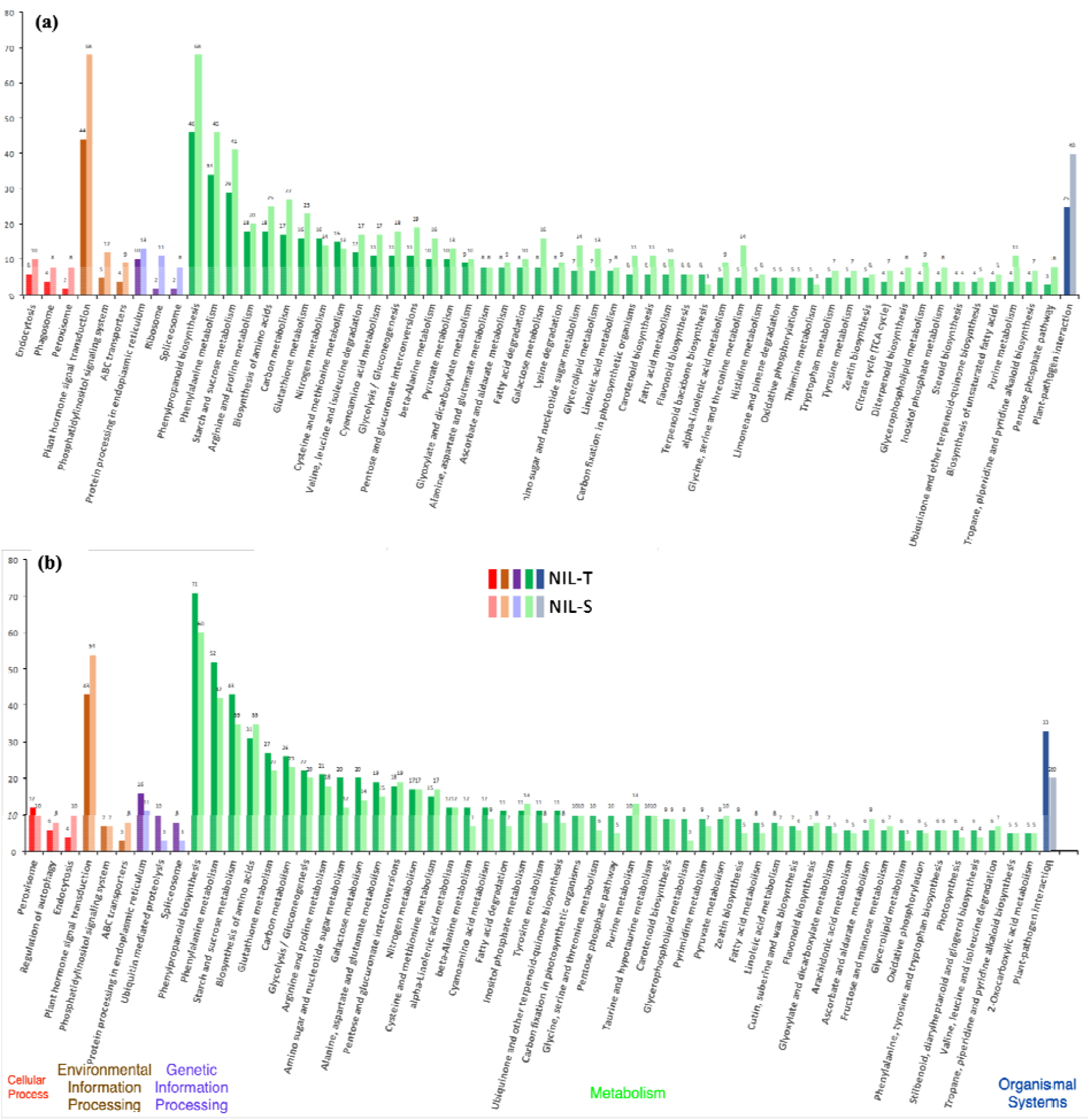
Significantly enriched KEGG (Kyoto Encyclopedia of Genes and Genomes) of all DEGs between salt-stressed and control samples in NIL-T and NIL-S soybeans. **a** 6h KEGG enrichment. **b** 3d KEGG enrichment. KEGG enrichment *P*-value cut-off ≤ 0.05.

After three days of salt stress (3d), 2844 and 2573 DEGs in NIL-T and NIL-S, respectively, were mapped to 57 pathways (corrected *p*-Value <0.05) (Fig. 4b). The metabolism category was also the most enriched after 3 days with similar pathways involved compared to 6 hours. After 3d salt-treatment, most of the significantly changed pathways included more DEGs in NIL-T compared to NIL-S, in particular in the pathway “Protein processing in endoplasmic reticulum” (16 to 11 DEGs), the cell organelle where GmSALT3 is localised (Guan *et al*., 2014). DEG enrichment in NIL-T was also higher in the pathways “Ubiquitin mediated proteolysis” (10 to 3 DEGs), “Phenylpropanoid biosynthesis” (71 to 60 DEGs), “Phenylalanine metabolism” (52 to 42 DEGs), “Starch and sucrose metabolism” (43 to 35 DEGs), and “Plant-pathogen interaction” (33 to 20 DEGs). It is important to note that all plants (NIL-T and NIL-S) were grown together and no plants showed signs of infection, suggesting that the genes categorised into this pathway also have a function in abiotic stress response.

In the “Plant-pathogen interaction” pathway, the putative CNGC (Cyclic Nucleotide-Gated ion Channel) 15-like gene, *Glyma*.*13G141000*, was significantly down-regulated in 3d T (Supplementary Fig. 5a; 5b), probable putative CNGC 20-like gene, *Glyma*.*09G168700*, was upregulated in 3d S. Several CaM (Calmodulin) and CML (Calmodulin-like) genes were down-regulated in 3d T. CDPK (Calcium-Dependent Protein Kinase 3, *Glyma*.*08G019700*) and Rboh (Respiratory Burst Oxidase Homolog protein F-like isoform 1, *Glyma*.*01G222700*) were up-regulated in 3d S. However, at 6h, another CNGC 20-like gene (*Glyma*.*16G218300*), homolog to the Arabidopsis putative Ca^2+^ channel that is proposed to regulate cytosolic Ca^2+^activity and expression of Rhob and CaM/CML (Demidchik *et al*., 2018), was significantly up-regulated in NIL-T (Supplementary Fig. 5d).

**Fig. 5.**
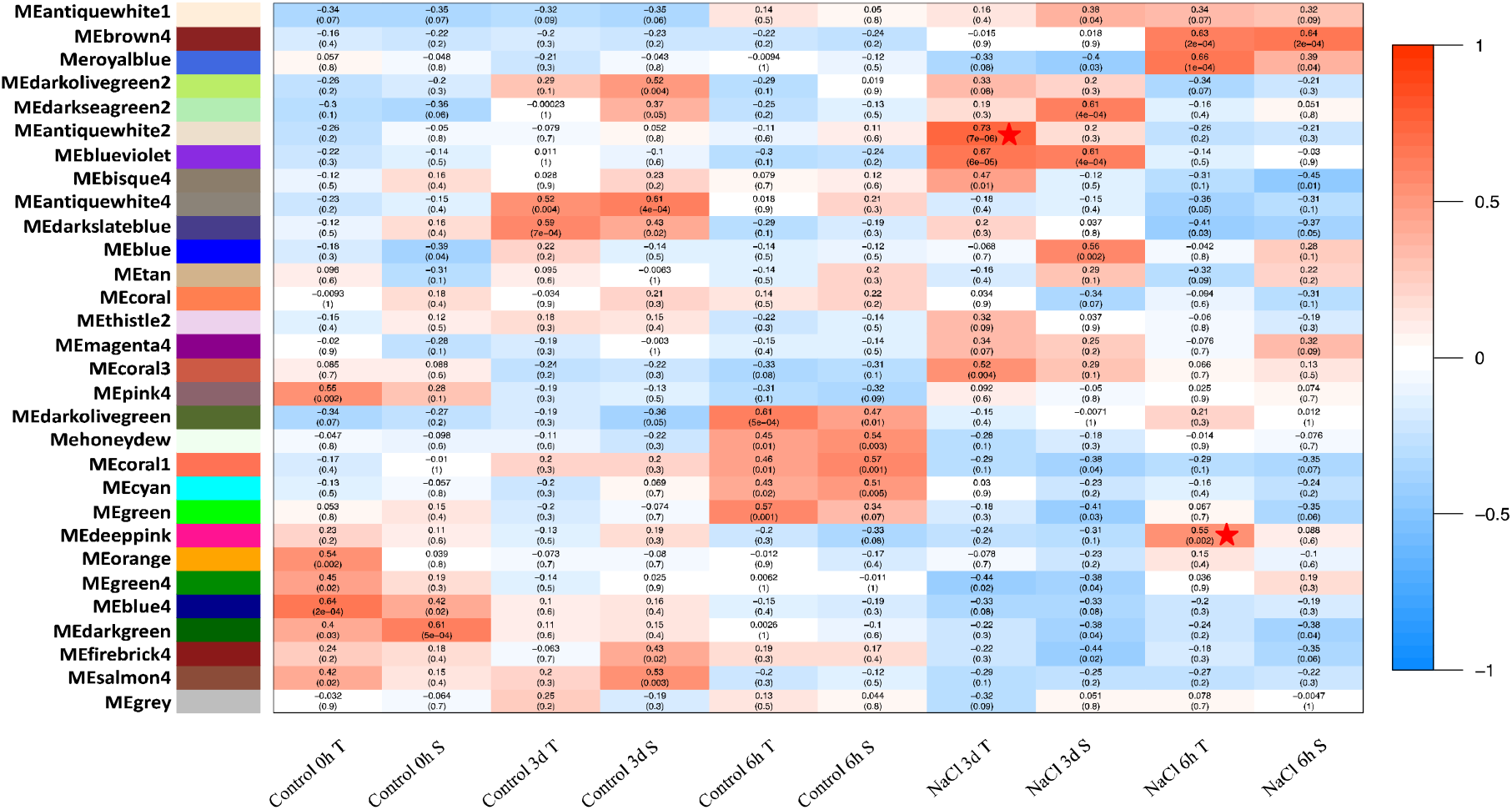
Module -Trait Relationships (MTRs) and corresponding *p*-values (within brackets) between the detected modules on the y-axis and groups (trait) on the x-axis. The MTRs are coloured based on their correlation: from red (strong positive correlation) to blue (strong negative correlation). Each group contains three biological replicates. Pentagram represents modules of interest.

### Weighted Gene Co-expression Network Analysis (WGCNA)

WGCNA identifies genes that have strongly correlated expression profiles, and those genes work cooperatively in related pathways to contribute to corresponding phenotypes. It is based on the concept that genes with strongly correlated expression profiles are clustered because those genes work associatively in related pathways, contributing to the resulting phenotype; in our case, establishing salinity tolerance after salt treatment. We applied the WGCNA to the normalized FPKM data (FPKM > 2) from all 30 RNA-seq libraries.

A co-expression network was constructed, and highly co-expressed genes were assigned to different colour-coded modules (Supplementary Fig. 6). Clusters were summarized by correlating the module’s eigengene (or an intromodular hub gene) to the sample traits (Langfelder and Horvath, 2008). Module -Trait Relationships (MTRs) were computed based on Pearson’s correlation between module and phenotypes, and then be used to screen modules for downstream analysis. Thirty clusters of highly co-expressed genes (modules) were detected and are shown in differently coloured module (Fig. 5). To investigate what genes show strongly correlated co-expression in NIL-T under saline conditions (6h and 3d), two modules of interest were selected for functional annotation based on their correlation value and corresponding *p*-Value. One module contains 79 genes that show strongly correlated co-expression in NIL-T after 6h salt treatment; this model is depicted here in a shade of pink (MEdeeppink). The second selected module, colour-coded here in a shade of white (MEantiquewhite2), contains 406 genes which expression profiles are highly correlated in NIL-T after 3d salt treatment.

**Fig. 6.**
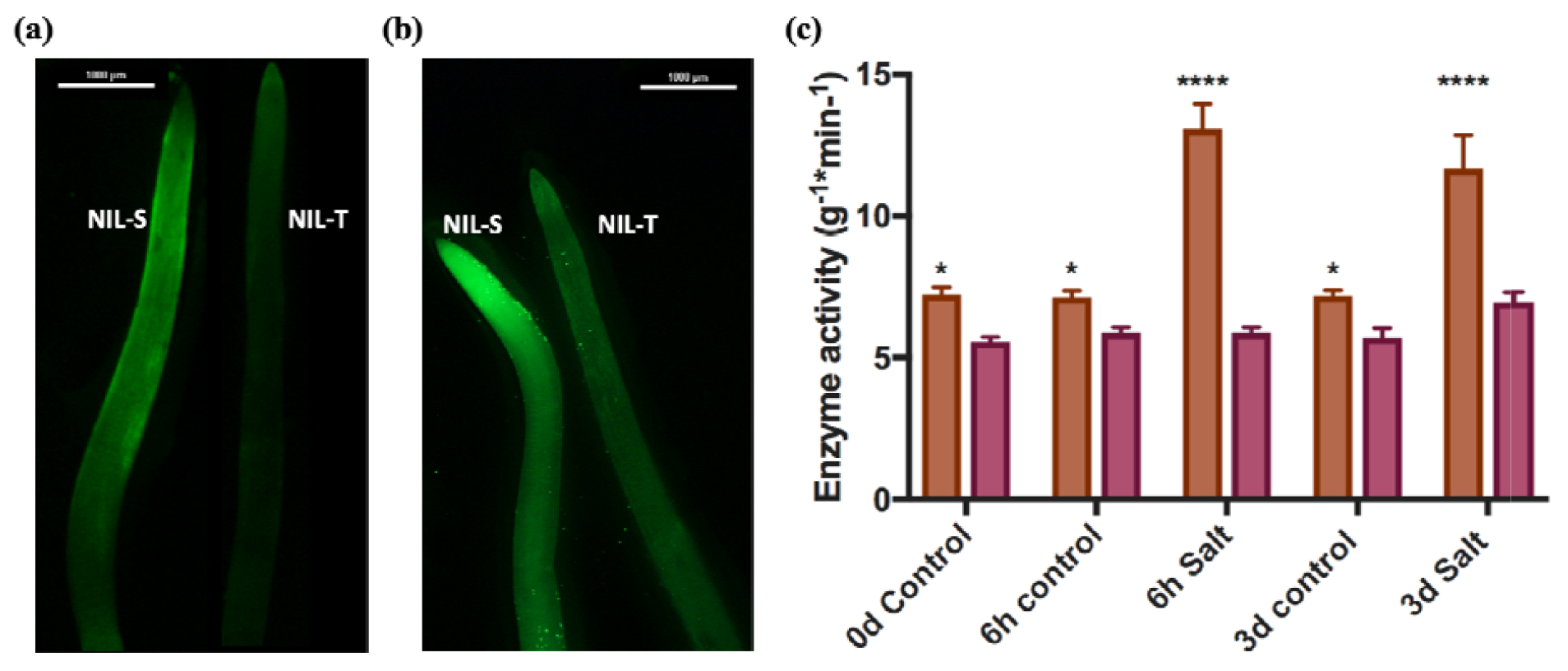
Relative ROS concentrations and enzyme activity of the superoxide anion (O_2_^-^) scavenger assay in soybean roots. NIL-T and NIL-S were labelled with 30 μM DCFDA for 10 min under control conditions (a) and with salt treatment for 3 days (b). Green fluorescence indicates the presence of ROS. (c) scavenging activity of the superoxide anion (O_2_^-^) of NIL-T (brown) and NIL-S (red) with or without salt treatment. Asterisk indicates a significant difference between NIL-T and NIL-S at *P <0.05 ****P<0.0001.

GO analysis of 79 genes in module MEdeeppink is shown in Table 5a; GO analysis of 406 genes in module MEantiquewhite2 is shown in Table 5b. Within the highly co-expressed 79 genes in NIL-T after 6h salt treatment (MEdeeppink module), Gene Ontology (GO) analysis indicates 12 of them are integral to plasma membrane (GO:0016021; Table 5a), and they encode a group of Casparian strip membrane proteins (Supplementary Table 1). Highly co-expressed 406 genes in NIL-T after 3d salt treatment (MEantiquewhite2 module) are enriched in GO: 0016021 (integral to membrane; 26 genes), which are related to intracellular vesicle trafficking and ion transport (Table 5b). Within those GOs, there are genes relevant to vesicle transport, such as genes encoding for coatomer subunits and transmembrane protein transporters; three potassium transporters including Glyma05g37270 (TRH1; tiny root hair 1), Glyma08g39860 (KUP6; potassium ion uptake transporter 6), and Glyma13g23960 (Supplementary Table 1).

**Table 5.**
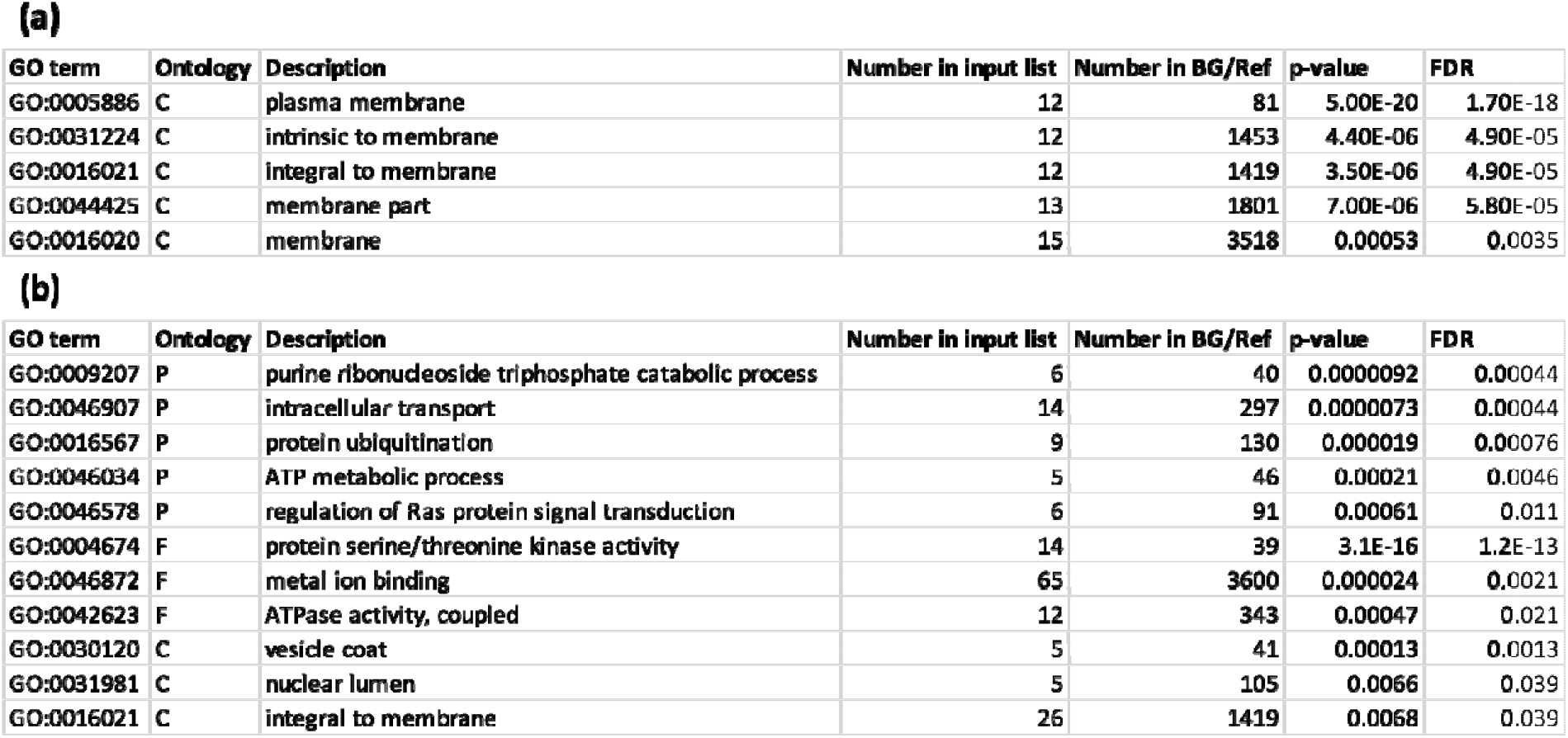
Significant Gene Ontology (GO) terms in MEdeeppink and MEantiquewhite2 modules. (a) Enriched GO terms in MEdeeppink module (NIL-T after 6h salt treatment); (b) Enriched GO terms in MEantiquewhite2 module (NIL-T after 3d salt treatment). C, Cellular Component; P, Biological Process; F, Molecular Function.

### RT-qPCR validation for RNA-seq results

In order to confirm the RNA-seq results, 10 DEGs were selected for RT-qPCR validation based on their RPKM transcript abundance. Importantly, this occurred on independently grown plants not used for RNAseq analysis but grown in identical conditions. RT-qPCR indicated that relative expression values (relative to housekeeping gene *GmUKN1*) of the selected DEGs were significantly correlated with their FPKM values (supplementary Fig. 7).

### ROS production and scavenger enzymes activity

Due to the enrichment of the “Oxidation reduction” gene ontology category in NIL-T plants under salt (Table 4), we decided to test if this translated into a difference in ROS generation or detoxification in NIL-T compared to NIL-S, again on independently grown plants. The ROS activity was measured in the roots of NIL-T and NIL-S after a 3-day treatment with or without salt treatment, using a fluorescent dye (Fig. 6a; 6b). We could detect a lower ROS activity in NIL-T compared to NIL-S roots, already in the control conditions, consistent with the expression of *GmSALT3* under control conditions. This difference strongly increased when comparing plants from saline conditions; ROS were strongly produced in NIL-S under salt treatment.

ROS are composed of many different molecules; we initially measured the concentration of the ROS H_2_O_2_. We could detect a higher H_2_O_2_ concentration in both lines under salt treatment compared to control; but no concentration differences between NIL-T and NIL-S in either control or salt treatment could be measured (Supplementary Fig. 9), suggesting that the increase in ROS is attributed to a different molecule. The antioxidant properties of the roots of NIL-T and NIL-S with or without salt treatment was then analysed using guaiacol. The antioxidant enzyme activity was assayed by measuring the absorbance at 470 nm due to the enzyme dependent oxidation of guaiacol by H_2_O_2_ (Pi *et al*., 2016). The scavenging enzyme activity of the superoxide anion was significantly higher in NIL-T compared to NIL-S under control and saline conditions (Fig. 6c).

## Discussion

Soybean cultivars harbouring *GmSALT3* are better able to modulate Na^+^, K^+^, and Cl^−^ content of shoots under salinity (Guan *et al*., 2014; Qi *et al*., 2014; Do *et al*., 2016; Liu *et al*., 2016; Qu *et al*., 2021), and they have a yield advantage under saline conditions compared to soybean varieties without a functional *GmSALT3* (Do *et al*., 2016, Liu *et al*., 2016). The GmSALT3 ion transport protein is ER-localised (Guan *et al*., 2014), suggesting a cellular function of GmSALT3 that is not directly connected to ion uptake or exclusion. We performed detailed and extended RNAseq analysis of NIL-T and NIL-S soybeans to decipher the cellular pathways and processes that connect *GmSALT3* to soybean salinity tolerance. We could identify several genes and biological pathways involved in responses to salt stress in *GmSALT3*-containing NIL-T plants, which gives an insight into how *GmSALT3* expression translates into a phenotype thorough modulating cell processes.

Illumina sequencing results showed that all the 30 constructed RNA-seq libraries were of sufficient quality for further analysis. On average the RNA-seq libraries had 26.8 million clean mapped reads, and reads were mapped to 53,625 annotated soybean genes with 353 additional transcripts not previously annotated – probably due to the germplasm being genetically divergent from the reference. Minimal DEGs could be detected between NIL-T and NIL-S at 0h, 6h, and 3d under control conditions (Fig. 2), this indicates that NIL-T and NIL-S roots had similar gene expression patterns under non-stressed conditions. In the three consistently up-regulated DEGs in NIL-S at all time points under control conditions, there was a gene encoding for a LOX (Linoleate 13S-lipoxygenase 3-1) (Fig. 2c). Lipoxygenases (LOXs) are involved in defence responses against biotic stresses, such as microbial pathogens and insect pests, in tomato (Hu *et al*., 2015), potato (Royo *et al*., 1999), maize (Gao *et al*., 2009), and rice (Zhou *et al*., 2009). The full length *GmSALT3* during soybean natural selection and domestication was lost several times independently (Guan *et al*., 2014), suggesting that plants with a non-functional GmSALT3 might have a benefit when grown under non-saline conditions. This benefit might be related to higher LOXs expression in soybean cultivars harbouring the sensitive alleles, which could be beneficial under certain biotic stresses. In potato, depletion of one specific LOX (LOX-H3) negatively affected their response to stress (Royo *et al*., 1999), but here in soybean, plants with the full length *GmSALT3* had lower LOX expression than with the non-functional GmSALT3. Therefore, this calls for further research to investigate the role of these LOX in soybean function and whether there is a link to GmSALT3 in response to abiotic stresses in soybean.

Comparing DEGs, the GO terms “Oxidation reduction (GO:0055114)” and “oxidoreductase activity (GO:0016491)” were consistently up-regulated in NIL-T at 6h and 3d of salt treatment, with a greater number of genes in this term up-regulated after 3 days, which indicates that NIL-T plants may have a greater capacity to detoxify ROS (Reactive oxygen species) than NIL-S plants. We could experimentally confirm this and show that more ROS is produced NIL-S compared to NIL-T (Fig. 6a; 6b) and NIL-T possessed a significantly higher scavenging enzyme activity of the superoxide anion (O_2_^−^) (Fig. 6c). Phenylalanine is the precursor for flavonoids, and flavonoids are utilised to scavenge and protect against ROS (Dastmalchi *et al*., 2016). The increase in gene expression related to phenylalanine synthesis seen in NIL-T (Fig 4a; 4b) is consistent with the observation that GmSALT3 is connected to an increase in general ROS detoxification capacity.

In all the listed up-regulated genes in NIL-T (Table 4), a group of Cytochrome P450 enzymes-coding genes were significantly more highly expressed in NIL-T in response to salt-treatment, especially *Glyma*.*13G173500*, which has the highest expression level and a fold change of 3.93. The cytochrome P450 (CYP) family is a large and essential protein superfamily in plants, and CYPs catalyse monooxygenation/hydroxylation reactions in primary and secondary metabolism pathways (Mizutani and Ohta, 2010). NaCl can induce conformational change of cytochrome P450 in animals (Yun *et al*., 1996; Oyekan *et al*., 1999), but this has not been investigated in plants. In soybean, there are 322 identified CYPs, with only few of them having been functionally characterised (Guttikonda *et al*., 2010). *GmCYP82A3* (*Glyma*.*13g068800*) (Yan *et al*., 2016) and *GmCYP51G1* (*Glyma*.*07g110900*) (Pi *et al*., 2016) were shown to be involved in plant tolerance stresses such as salinity and drought. The two P450 containing isoflavone synthases (IFS) *GmIFS1* (*Glyma*.*07G202300*) and *GmIFS2* (*GmCYP93C1* = *Glyma*.*13G173500*) are both anchored in the ER, the same organelle where GmSALT3 is localised. *GmIFS2* is a DEG identified in our study.

Expression of *GmIFS1* was previously shown to be induced by salt stress, and has been linked to improving salinity tolerance through the accumulation of isoflavone content (Jia *et al*., 2017). We found its homologue, *GmIFS2* (*Glyma*.*13G173500)*, had a fold change of 3.93 in NIL-T in response to salt stress (Table 4). Combined, this suggests that the function of GmSALT3 could be important for the synthesis of isoflavones, and this may contribute to salt tolerance.

ROS production has been linked to Ca^2+^ influx into cells and specifically the activity of CNGCs (Ma *et al*., 2009). Ca^2+^ and ROS act as signalling molecules when plant roots are suddenly exposed to salt (Byrt *et al*., 2018). Gene expression related to proposed Ca^2+^ signalling elements are different between NIL-T and NIL-S. As a result, our analysis suggested that the “Plant-pathogen interaction” pathway could be highly activated in NIL-S, which may lead to higher ROS production and hypersensitive responses (Supplementary Fig. 5a; 5c). Under 6h salt treatment, several CNGCs were also significantly up-regulated in NIL-S, which may relate to activation of different signalling cascades (Supplementary Fig. 5d). Several CaM (Calmodulin) and CML (Calmodulin-like) genes are also down-regulated in 3d T (Supplementary Fig. 5b), down-regulation of CaM and CML may has been proposed to restrict gaseous reactive oxygen species (ROS) production in plants (Ma and Berkowitz, 2011).

The WGCNA analysis revealed that genes related to Casparian strip (CS) development were upregulated in NIL-T within 6 hours of salt treatment. CSs play an important role in regulating nutrient uptake and salt stress tolerance, as they form the diffusion barrier for ions into the stele (Chen *et al*., 2011; Roppolo *et al*., 2011). CS localised membrane proteins (CASPs) were shown to mediate CS formation in Arabidopsis, by guiding the local lignin deposition during cell wall modification (Roppolo *et al*., 2011). Upon immobilisation, CASPs recruit PER64 (Peroxidase 64), RBOHF (Respiratory Burst Oxidase Homolog F), ESB1 (Enhanced Suberin 1), and dirigent proteins, to establish the lignin polymerization system (Hosmani et al., 2013; Kamiya et al., 2015; Lee et al., 2013). Our analysis identified a module of genes upregulated under salt exclusively in the NIL-T (MKdeeppink module). This module contains 8 dirigent genes, peroxidase 64 (Glyma.14g221400) as well as CASPs. The combination of genes in this model suggests that NIL-T plants might more efficiently induce an earlier formation of the endodermal diffusion barrier as well as the formation of an exodermal diffusion barrier, in response to salt stress. We stained lignin and suberin in NIL-T and NIL-S roots with ClearSee and Auramine O, following the protocol developed by Ursache *et al*. (2018), in order to compare the CS development. Unfortunately, we could only observe autofluorescence in both NIL-T and NIL-S root differentiation zone that close to the root tips using confocal microscopy. So the link between GmSALT3 and CS formation needs further validation.

Lastly, CHX transporters (of which GmSALT3 is a member) were shown to be associated with the endocytic and exocytic pathways in dynamic endomembrane system (Chanroj *et al*., 2012; Padmanaban *et al*., 2007). Our results and the previously proposed vesicle trafficking function of CHXs (Chanroj *et al*., 2012), suggests that GmSALT3 might be important for establishing optimal conditions in the ER lumen, which could subsequently impact the secretion of compounds such as signalling molecules, ions, and soluble proteins.

To summarize, our findings suggest that the GmSALT3 containing NIL-T has a higher salinity tolerance because NIL-T roots are more effective in scavenging ROS, which helps to prevent cellular damage. Our results suggest that the improved ROS detoxification mechanisms might be connected to ER-localised flavonoid biosynthesis enzymes. Additionally, our study has identified that Ca^2+^ signalling, vesicle trafficking and Casparian strip development might be modulated by GmSALT3, and are therefore areas for further study.

## Supplementary data

**Supplementary Fig. 1 GO term analysis of unique DEGs under salt treatment in NIL-T roots after 6h 200 mM NaCl treatment**.

**Supplementary Fig. 2 GO term analysis of unique DEGs under salt treatment in NIL-S roots after 6h 200 mM NaCl treatment**.

**Supplementary Fig. 3 GO term analysis of unique DEGs under salt treatment in NIL-T roots after 3d 200 mM NaCl treatment**.

**Supplementary Fig. 4 GO term analysis of unique DEGs under salt treatment in NIL-S roots after 3d 200 mM NaCl treatment**.

**Supplementary Fig. 5. DEGs between salt-treated and control samples of NIL-T and NIL-S in plant-pathogen interaction KEGG pathway at 6h and 3d**.

**Supplementary Fig. 6 Gene dendrogram showing the co-expression modules defined by the WGCNA labelled by colours**.

**Supplementary Fig. 7 qPCR validation for RNA-seq results of selected genes. Supplementary Fig. 8 *Glyma*.*13G173500* (*GmIFS2*) gene expression (FPKM) in all samples**.

**Supplementary Fig. 9 H**_**2**_**O**_**2**_ **concentration measurement in soybean roots**.

**Supplementary Table 1 gene annotations in each GO (attached as an Excel file)**.

## Acknowledgements

This work was funded by the Natural Science Foundation of China, grant number 31830066 (RG); Scientific Innovation Project of Chinese Academy of Agricultural Sciences (L-J Q). YQ and SH were supported by ARC Centre of Excellence funding awarded to MG (CE140100008). ARC Fellowships supported MG (FT130100709) and SW (DE160100804). We thank Na Sai and Dr. Miriam Schreiber for assisting in WGCNA analysis.

## Author contributions

MG, YQ and RG designed experiments, with input from SW and LQ. YQ and RG analysed the data. LQ and MG directed the project. OB and JW contributed to RNA-sequencing. LY performed plant samples harvest for RNA-sequencing. RD contributed to RNA-sequencing analysis. YQ, MG and RG wrote the paper. SW and LQ assisted in editing the final version of the manuscript. All authors commented on the manuscript.

## Data availability statement

The data supporting the findings of this study are available from the corresponding author, Matthew Gilliham, upon request.

